# Extreme niche partitioning promotes a remarkably high diversity of soil microbiomes across eastern Antarctica

**DOI:** 10.1101/559666

**Authors:** Eden Zhang, Loïc M. Thibaut, Aleks Terauds, Sinyin Wong, Josie van Dorst, Mark M. Tanaka, Belinda C. Ferrari

**Affiliations:** School of Biotechnology and Biomolecular Sciences, University of New South Wales, Sydney, 2052, Australia.; Australian Antarctic Division, Department of Environment, Antarctic Conservation and Management, 203 Channel Highway, Kingston, TAS, Australia, 7050.

**Keywords:** Antarctica, Soil Microbiome, Species Abundance Distribution, Bacteria, Eukarya, Archaea

## Abstract

Terrestrial Antarctica, a predominantly microbial realm, encompasses some of the most unique environments on Earth where resident soil microbiota play key roles in the sustainability and evolution of the ecosystem. Yet the fundamental ecological processes that govern the assemblage of these natural communities remain unclear. Here, we combined multivariate analyses, co-occurrence networks and fitted species abundance distributions of amplicon sequencing data to disentangle community assemblage patterns of polar soil microbiomes across two ice-free deserts (Windmill Islands and Vestfold Hills) situated along the coastline of eastern Antarctica. Our findings report that communities were predominantly structured by non-neutral processes, with niche partitioning being particularly strong for bacterial communities at the Windmill Islands. In contrast, both eukaryotic and archaeal communities exhibited multimodal distributions, indicating the potential emergence of neutrality. Between the three microbial domains, polar soil bacterial communities consistently demonstrated the greatest taxonomic diversity, estimated richness, network connectivity and linear response to contemporary environmental soil parameters. We propose that reduced niche overlap promotes greater phylogenetic diversity enabling more bacterial species to co-exist and essentially thrive under adversity. However, irrespective of overall relative abundance, consistent and robust associations between co-existing community members from all three domains of life highlights the key roles that diverse taxa play in ecosystem dynamics.

**Significance:** In the face of a warming Antarctica, contemporary dynamics between polar soil microbial communities will inevitably change due to the climate-induced expansion of new ice-free areas. Increasing concern about disturbance and rapid biodiversity loss has intensified the need to better understand microbial community structure and function in high-latitude soils. We have taken an integrated approach to elucidate domain-level assemblage patterns across east Antarctic soil microbiomes. These assemblage patterns will be available to feed into policy management and conservation planning frameworks to potentially mitigate future biodiversity loss.

## Introduction

East Antarctica constitutes up to two-thirds of the Antarctic continent and is home to some of the oldest, coldest soils on Earth (Cary et al., 2010). Aside from isolated pockets of ice-free areas, its sheer bulk is typically covered by a thick layer of ice (Terauds et al., 2017). The Windmill Islands, an ice-free region situated near Casey research station, is comprised of five major peninsulas and a number of rock-strewn islands. Approximately 1400km north lies the Vestfold Hills, a large expanse of low-lying hilly country deeply indented with sea-inlets and snowmelt lakes (O’Brien et al., 2015). These diverse edaphic habitats are a legacy of varied geological and glaciological histories (Anderson et al., 2002).

Both contemporary and historical conditions are believed to drive the current biogeography of soil microbiota across the Antarctic continent (Chown & Convey 2007; Convey et al., 2015; Cowan et al., 2014; Ferrari et al., 2016; Terauds et al., 2012). Largely dictated by microclimate and soil age, abiotic factors such as water, energy and nutrient availability have been reported to notably influence Antarctic species distributions and life histories (Aislabie et al., 2008; Cary et al., 2010; Convey et al., 2014; Siciliano et al., 2014; Terauds et al., 2012). These properties can co-vary with local lithology, pedology and geographical position, leading to a myriad of edaphic niches (Chong et al., 2012). In turn, their microbial occupants are fundamental to establishing and maintaining core ecosystem processes, occasionally involving unique taxa and novel functional traits (Benaud et al., 2019; Cary et al., 2010; Chan et al., 2013; Ji et al., 2017).

Throughout terrestrial Antarctica, resources are scarce and physiochemical gradients steep (Convey et al., 2014). It is therefore hypothesised that variation in the capacity of microbes to access and utilise resources, as well as tolerate stress, is contributing significantly to the structuring of these microbial assemblages inhabiting cold desert soils. But, the ability to disentangle the basis of microbial community assembly, specifically niche-neutral processes in cold regions has been limited by the small number, and the depth of studies available (Cowan et al., 2014). Furthermore, the majority of relevant studies have solely been focused on a portion of the microbiome, the bacterial community.

Relatively few eukaryotic and archaeal-specific phylotypic surveys have been reported for terrestrial Antarctic environments (Cowan et al., 2014). As a result, the ecological roles of eukaryotes and archaea in cold edaphic habitats remain ambiguous (Pointing et al., 2009; Rao et al., 2012; Richter et al., 2015). Available studies report significantly lower fungal and archaeal diversities within arid-to-hyperarid soil ecosystems compared to their bacterial counterparts (Cowan et al., 2014; Ferrari et al., 2016). However, lower diversity and abundance does not necessarily equate to a diminished ecological role. In mixed soil communities, it is often not the most productive members that dominate as relative abundance is often determined by adaptations to the abiotic and biotic components of the environment (Bell et al., 2013). As such, it is likely that all three microbial domains are collectively responsible for the sustainability and evolution of the polar soil microbiome (Faust & Raes 2012; Fierer 2017). Therefore, in order to approach an integrated understanding of the basic ecological mechanisms behind community assemblage patterns within such a severely limiting environment, it is important to jointly consider their bacterial, eukaryotic and archaeal components together,

In this study, we compiled bacterial 16S, eukaryotic 18S and archaeal 16S rRNA amplicon sequencing data from over 800 polar soil samples spanning nine east Antarctic sites between the Windmill Islands and Vestfold Hills. By taking a multivariate, exploratory network and modelling approach using poisson-lognormal (PLN) and negative binomial (NB) fitted species abundance distributions (SADs), we aim to determine whether classic niche-based or neutral mechanisms best explain the assemblage patterns of microbial communities across our east Antarctic soil biomes.

## Results

### Amplicon sequencing yield and coverage

We recovered a total of 60, 495, 244 high-quality bacterial 16S rRNA gene sequences, which clustered down into 36, 251 operational taxonomic units (OTUs) at 97% identity cut-off. Our eukaryotic and archaeal runs yielded a total of 1, 299, 519 18S rRNA and 13, 373, 072 16S rRNA gene sequences after read-quality filtering, which respectively clustered at 97% into 1511 and 589 OTUs (Table S1). Subsampled rarefaction curves of the pooled data revealed that bacterial, eukaryotic and archaeal richness approached asymptote at each site (Fig. S1).

### Biodiversity of the east Antarctic polar soil microbiome

At 97% identity, OTUs were classified into 63 bacterial, 27 eukaryotic and three archaeal phyla. Distributions of phylum abundances for all three domains were uneven as the majority of sites were dominated by a handful of taxa (Fig. 2). Overall, our soil bacterial communities were predominantly comprised of the metabolically diverse *Actinobacteria* (30.5%) and *Proteobacteria* (14.6%). *Bacteroidetes* were more prevalent at the Vestfold Hills (24.9%) due to the higher salinity levels visible as salt crystal encrustations in this region. *Chloroflexi* (17.8%) and *Acidobacteria* (13.6%) were present in greater relative abundances throughout the Windmill Islands. With the exception of Browning Peninsula (BP=10.9%), Herring Island (HI=3.1%) and Rookery Lake (RL=4.2%), *Cyanobacteria* abundance was relatively low across all sites. At Mitchell Peninsula (MP) and Robinson Ridge (RR), rare candidate phyla namely *Candidatus Eremiobactereota (WPS-2)* and *Candidatus Dormibactereota (AD3)* were in significantly higher relative abundances (>4.6%) than other sites. At lower taxonomic levels, bacterial sequences classified into 169 classes, with members largely belonging to *Flavobacteria* (10.9%) and *Actinobacteria* (9.0%) followed by similar proportions (~6.0%) of *Thermoleophilia, Chloracidobacteria, Gamma-proteobacteria* and *Alpha-proteobacteria* (Fig. S2). As taxonomic levels decreased further, the number of unclassified bacterial sequences substantially increased (order=12.1%, family=31.5% and genus=61.0%).

**Figure 1.**
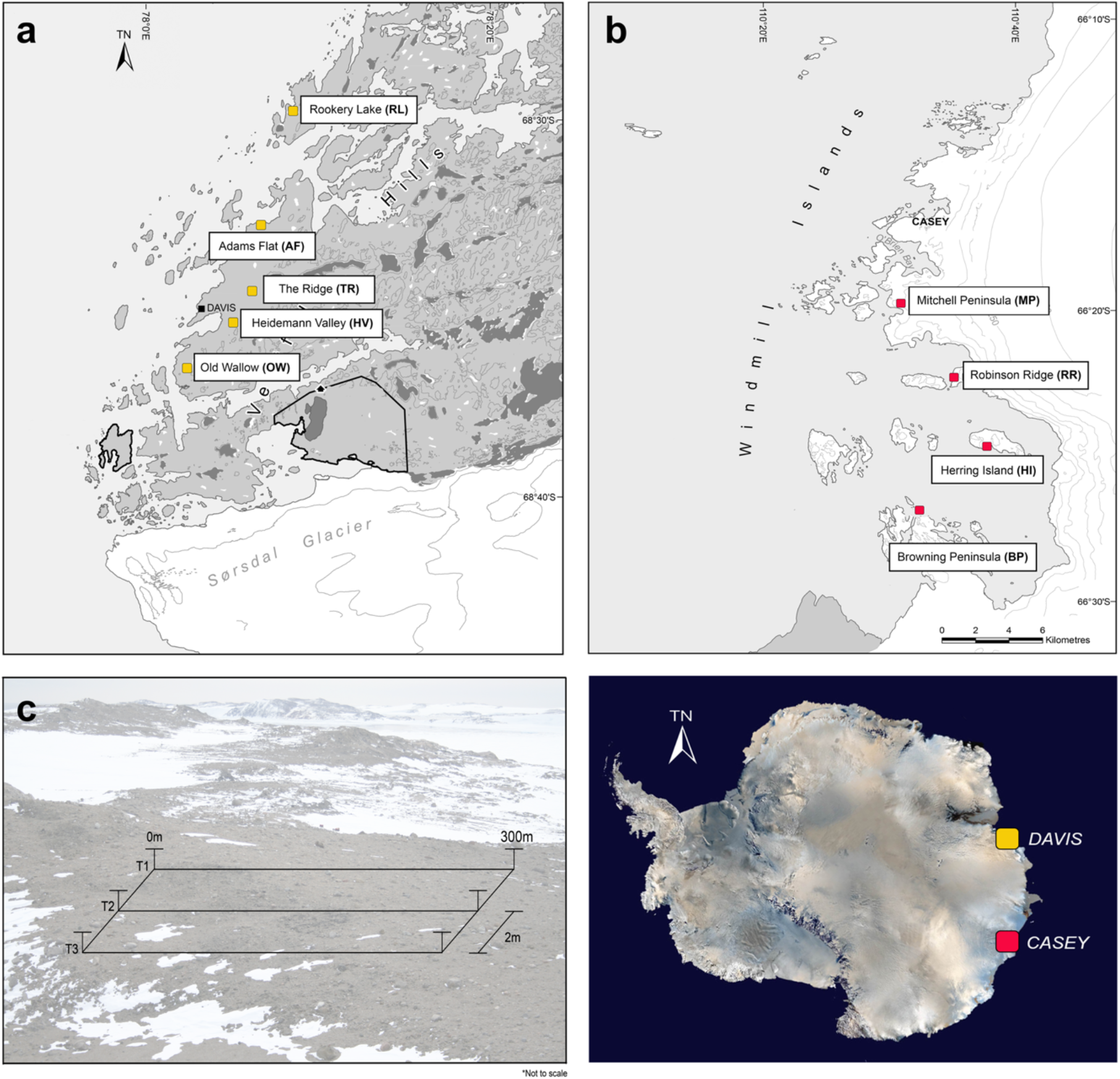
Map of the nine study areas across the (a) Vestfold Hills (AAD map catalogue No. 14499) and (b) Windmill Islands (No. 14179) region of Eastern Antarctica, showing approximate sampling locations and (c) geospatial transect design. At each site, soil samples (n=93) were taken at the following distance points along each transect: 0, 0.1, 0.2, 0.5, 1, 2, 5, 10, 20, 50, <u>100</u>, 100.1, 100.2, 100.5, 101, <u>102</u>, 105, 110, 120, 150, <u>200</u>, 200.1, 200.2, 200.5, 201, <u>202</u>, 205, 210, 220, 250 and 300m. Where <u>underlined</u> distance points refer to a subsample (n=18) submitted for amplicon sequencing of eukaryotic (18S rRNA) and archaeal (16S rRNA) communities.

**Figure 2.**
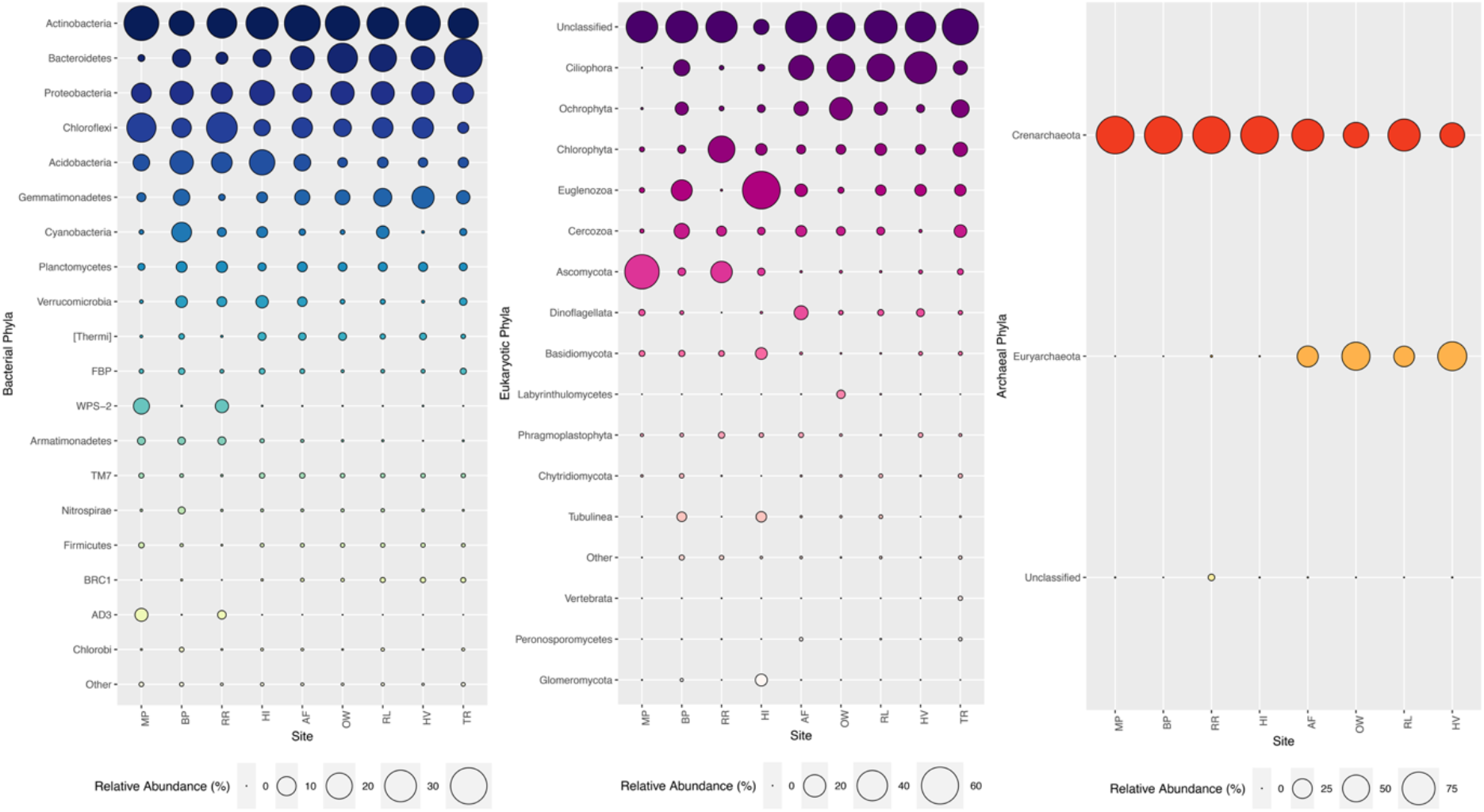
Bubbleplots of relative abundance (%) per site of phyla-level composition of OTUs (97% cut-off), based on bacterial 16S (mean=490bp), eukaryotic 18S (mean=125bp) and archaeal 16S (mean=470bp) SSU rRNA sequences representing >0.001% of all normalised OTUs sorted by decreasing relative abundance. Greatest phylogenetic diversity is exhibited by bacteria followed by eukarya then archaea. Across all three domains, distribution of phyla abundances is generally uneven as a handful of taxa tend to dominate but strong compositional differences are apparent between the Windmill Islands and Vestfold Hills regions.

For eukaryotes, 18S rRNA sequences fell into six supergroups consisting of unclassified (46.9%), *Chromalveolata* (i.e. *Ciliophora* and *Dinoflagellata* =20.6%), *Archaeplastida* (i.e. *Ochrophyta, Chlorophyta* and *Phragmoplastophyta=17.8%), Excavata* (i.e. *Euglenozoa* =5.4%), *Opisthokonta* (i.e. *Ascomycota, Basidiomycota, Labyrinthulomycetes, Chytridiomycota, Vertebrata, Peronosporomycetes* and *Glomermomycota* =4. 6%) and *Amoebozoa* (i.e. *Cercozoa* and *Tublinea* =4.4%). Fungal diversity contributed a fairly small proportion (10.5%) to the total relative abundance of our eukaryotic soil communities, except at MP and RR. Unclassified eukarya remained dominant across all taxonomic levels, with moderately higher relative abundance observed throughout the Vestfold Hills (61.3%) than the Windmill Island sites (38.8%), particularly at The Ridge (TR).

Archaeal diversity was mainly distributed within the *Crenarchaeota* phylum (84.54%), whilst members of *Euryarchaeota* (15.0%) were exclusive to the Vestfold Hills. In addition, an unusually high proportion (2.3%) of unclassified archaea was observed at RR. At lower taxonomic levels, archaeal sequences belonged to six main families consisting of *Nitrososphaeraceae* (84.5%) and *Halobacteriaceae* (15.0%), followed by unclassified, *SAGMA-X, Cenarchaeaceae* and *TMEG* families that collectively accounting for 0.01% of total relative archaeal abundance.

### Domain-level biotic interactions

Non-metric multidimensional scaling (NMDS) ordination of microbial OTU communities and corresponding environmental metadata revealed that samples were conserved between sites and broadly by geographic region (Fig. S3). bacterial communities exhibited the greatest overall species richness based on Chao1 estimates (Fig.3), particularly at the Windmill Islands (observed mean=1341.9, estimated mean=2270.1). In contrast, greater eukaryotic richness was observed throughout the Vestfold Hills (observed mean=56.1, estimated mean=132.3). archaeal communities exhibited the lowest overall species richness (observed mean=35.9, estimated mean=50.9), with RR being an exception (observed mean=94.6, estimated mean=106.4). Pearson’s correlations between domain-level pooled Chao1 richness estimates revealed weak but significant (*P*<0.05) negative relationships of bacterial communities against both eukaryotic (*R*=−0.23, *P*=0.0034) and archaeal (*R*=−0.17, *P*=0.045) communities. However, no significant correlation was found between eukaryotic and archaeal richness (*R*=0.039, *P*=0.64). Networks displaying the co-occurrence of OTUs offered new insights into the polar soil microbiome through the sharing of niche spaces or potential interactions between co-existing taxa at the domain level (Fig. 5). The resulting network for the Vestfold Hills consisted of 43 nodes (clustering coefficient=0.214) and 44 edges (average no. of neighbours=2.047, characteristic path length=3.247) across 8 connected components with a network diameter of seven edges (Table S2). Whereas, the resulting Windmill Islands network consisted of 58 nodes (clustering coefficient=0.448) and 201 edges (average no. of neighbours=6.931, characteristic path length=2.377) across three connected components with a network diameter of six edges (Table S2). Overall, microorganisms present within our soil microbial networks tended to co-occur more than expected by chance (*P*<0.001).

**Figure 3.**
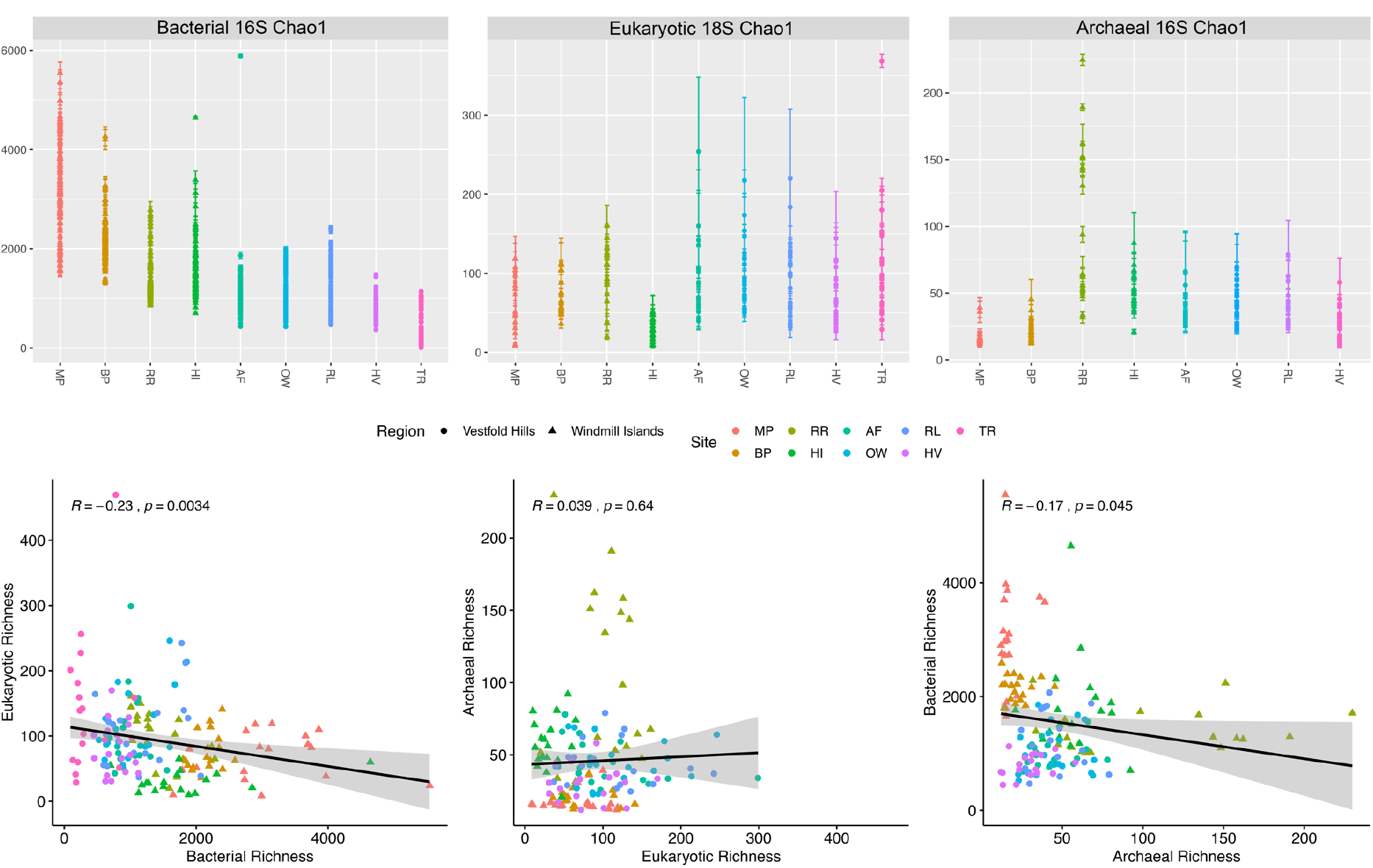
Chao1 richness estimates and correlations between our soil bacterial, eukaryotic and archaeal communities coloured by site. Our polar soil bacterial communities demonstrated highest overall species richness estimates, particularly throughout the Windmill Islands region. Significant (*P*<0.05) negative correlations were detected between estimated bacterial species richness against the other two microbial domains.

### Linear correlations between species richness and selected physiochemical soil factors

bacterial, eukaryotic and archaeal communities within these polar soils exhibited different response patterns when correlated against the selected physiochemical variables (Table 1). Out of the 52 variables tested, bacterial richness demonstrated the highest total number of significant correlations (n=26, *P*<0.05), the strongest relationships were observed for pH (*R*=0.63), SiO_2_ (*R*=−0.64), Al_2_O_3_ (*R*=0.68) and gravel (*R*=0.61). bacterial species richness was largely found to be negatively correlated against nutrient availability, water extractable ions and oxide levels (*P*<0.05). Whereas, positive associations were generally observed with particle size (*P*<0.05). In contrast, fewer significant correlations were observed for both eukaryotic (n=15, *P*<0.05) and archaeal (n=13, *P*<0.05) richness against the selected soil parameters, most of which comprised weak-to-moderate strength correlations. Shared correlations (n=15, *P*<0.05) between eukarya and bacteria inversely selected for richness such as particle size and oxide levels. Whereas, archaea were uniquely correlated to some variables of total carbon (TC, *R*=0.18, *P*<0.05), total nitrogen (TN, *R*=0.24, *P*<0.05) and Iron (Fe) content (*R*=0.49, *P*<0.05).

**Table 1.**
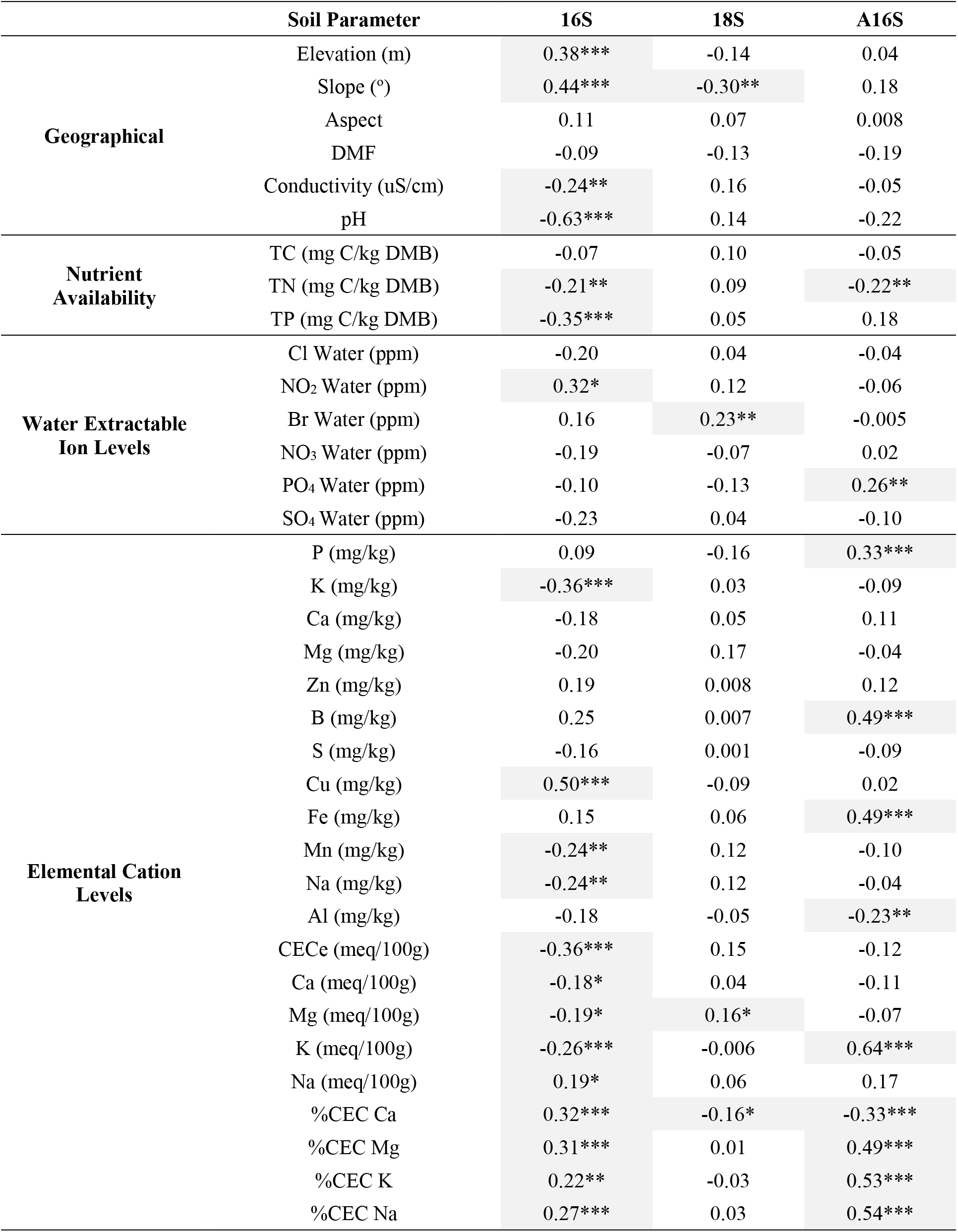
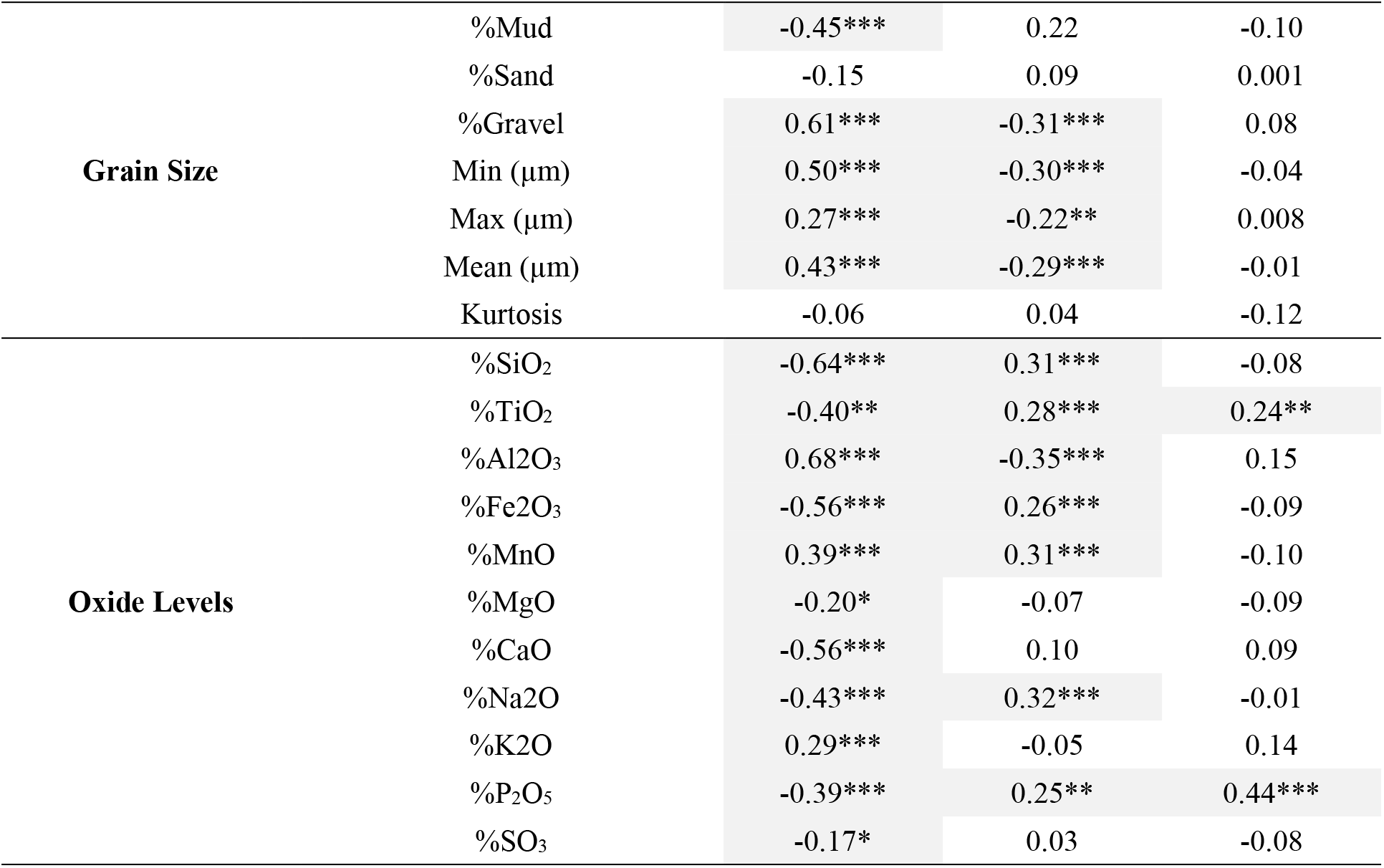
Pearson’s correlations between microbial richness estimates and selected physiochemical soil parameters (where significant correlations are shaded in grey and * = p < 0.05, ** = p < 0.01 and *** = p < 0.001).

### The niche-neutral debate

Overall, species abundances were better approximated by poisson-lognormal (PLN) distributions than the negative binomial (NB) distribution as these communities are substantially more heterogenous (Fig. 5). All distributions were characterised by highly left-skewed patterns, emphasising the disparity between rare and common species, which is a feature often associated with ecosystems subjected to periodic disturbance such as freeze-thaw cycles. Bacterial communities often lacked an internal mode and demonstrated the best PLN-fit, particularly at the Windmill Islands where there was an excess of rare species and species exhibiting intermediate abundances. In contrast, eukaryotic and archaeal communities demonstrated weaker PLN-fits with transient multimodal distributions suggesting the emergence of neutrality. Interestingly, these trends were also consistently observed at the site level (Fig. S4).

## Discussion

Remarkably strong niche partitioning was found to be driving the establishment and maintenance of contemporary microbial communities in the arid-to-hyperarid east Antarctic soils analysed here, particularly bacterial communities (Fig. 5 and S4). In addition, neutral processes played larger than expected role within the relatively species-poor eukaryotic and archaeal communities. This outcome supports the shift towards a more unified concept of biodiversity where deterministic-niche and stochastic-neutral processes are not mutually exclusive (Dini-Andreote et al., 2014; Dumbrell et al., 2010; Scheffer et al., 2018; Vellend 2010). Within such a severely limiting yet dynamic environment like Antarctica, niche differences or similarities amongst species would be expected to promote long-term species coexistence (Scheffer et al., 2018; Verbeck 2011).

Both PLN and NB distributions have been proven to be accurate in predicting both niche and simulated neutral models, especially when datasets contain a high number of abundance values and are pooled at the meso- or regional scale (Connolly et al., 2013). SADs are providing valuable insight into less visible aspects of community assembly such as competition and predation, for example when two species occur together, yet never at high densities (Verbeck 2011). However, it should be noted that an important limitation of current SAD sampling theories is the omission of species identity (Alonso et al., 2008). Nonetheless, comparisons of our PLN- and NB-fitted SAD curves here offered robust visualisations of non-neutrality signatures on some of the most pristine and undisturbed natural communities on Earth (Terauds et al., 2012).

Very strong niche partitioning are involved in the structuring of our polar soil bacterial communities (Fig. 5 and S4; Table 1). It was particularly evident for bacterial communities at the Windmill Islands where environmental gradients were more pronounced (Fig S3 and Table S3). As theorised, reduced niche overlap may result in weaker interspecific competition that aids coexistence within species-rich communities (Finke & Snyder 2008), a feature that promotes greater biodiversity and resource exploitation by the relatively species-rich bacterial communities existing under adverse conditions (Fig. 2 and 3). In contrast, species-poor communities inhabiting more homogenous environments, neutral dynamics may dominate as dispersal limitation or longer lifespans may prevent competitive exclusion (Verbeck 2011).

At both regional and local scales, apparent multimodality of the relatively species-poor eukaryotic and archaeal communities suggest that neutral processes play a larger role in ecosystem processes than expected, particularly at the Vestfold Hills (Fig. S5 and S4). While there is no current consensus about what drives SAD shape variation, multimodality is rarely reported and its implications not well understood (Antão et al., 2017). A number of studies argue that multimodality occurs quite frequently in nature, and as such it is indeed a characteristic of ecological communities (Antão et al., 2017; Dornelas & Connolly 2008; Matthews et al., 2015; Vergnon et al., 2012). Emergent neutrality is a hypothesis put forth that explains multimodal SADs, in which self-organised patterns of functionally similar species that coexist within an ecological niche (Holt 2006; Vergnon et al., 2012). These patterns are likely to be transient (Scheffer et al., 2018). However, at very high similarities the displacement of the weaker competitor becomes exceedingly slow (Scheffer et al., 2018). Such as phenomenon was clearly reflected in our polar soil archaeal communities, where multimodality was observed across both regions (Fig. 5), with members belonging to the functionally important *Crenarchaeota* phylum (Fig. 2), present >85% abundances throughout the entire dataset. *Nitrosospharae* (Fig. S2), a genus of chemotrophic ammonia oxidisers, implicated in nitrogen cycling in nutrient-limited Antarctic soils, also dominated (Tourna et al., 2011). Interestingly, draft genomes of *Thaumarchaeota* recovered from our soils reported the presence of ammonia monooxygenase (Ji et al., 2017), the first enzyme in the pathway for nitrification (Pester et al., 2012).

In support of previous findings in arid soil environments (Cowan et al., 2014), this survey of east Antarctic soils reveal that while bacterial diversity is high, both eukaryotic and archaeal diversities are relatively low (Fig. 2 and 3). This disparity likely reflects differing ecological roles and life history strategies (De la Riva et al., 2018). More importantly, their respective activities, particularly those forming metabolic alliances with or competing against other species, are critical to the formation of functional microbial communities within these harsh environments (Aller et al., 2008; Bell et al., 2013). Our soil microbial networks were comprised of highly modular structures, consisting of co-occurring OTUS from all three domains (Fig. 4). Thereby, contributing to the integrative concept that bacteria, eukarya and archaea are likely altogether responsible for structuring and maintaining the polar soil microbiome (Bahram et al., 2018).

**Figure 4.**
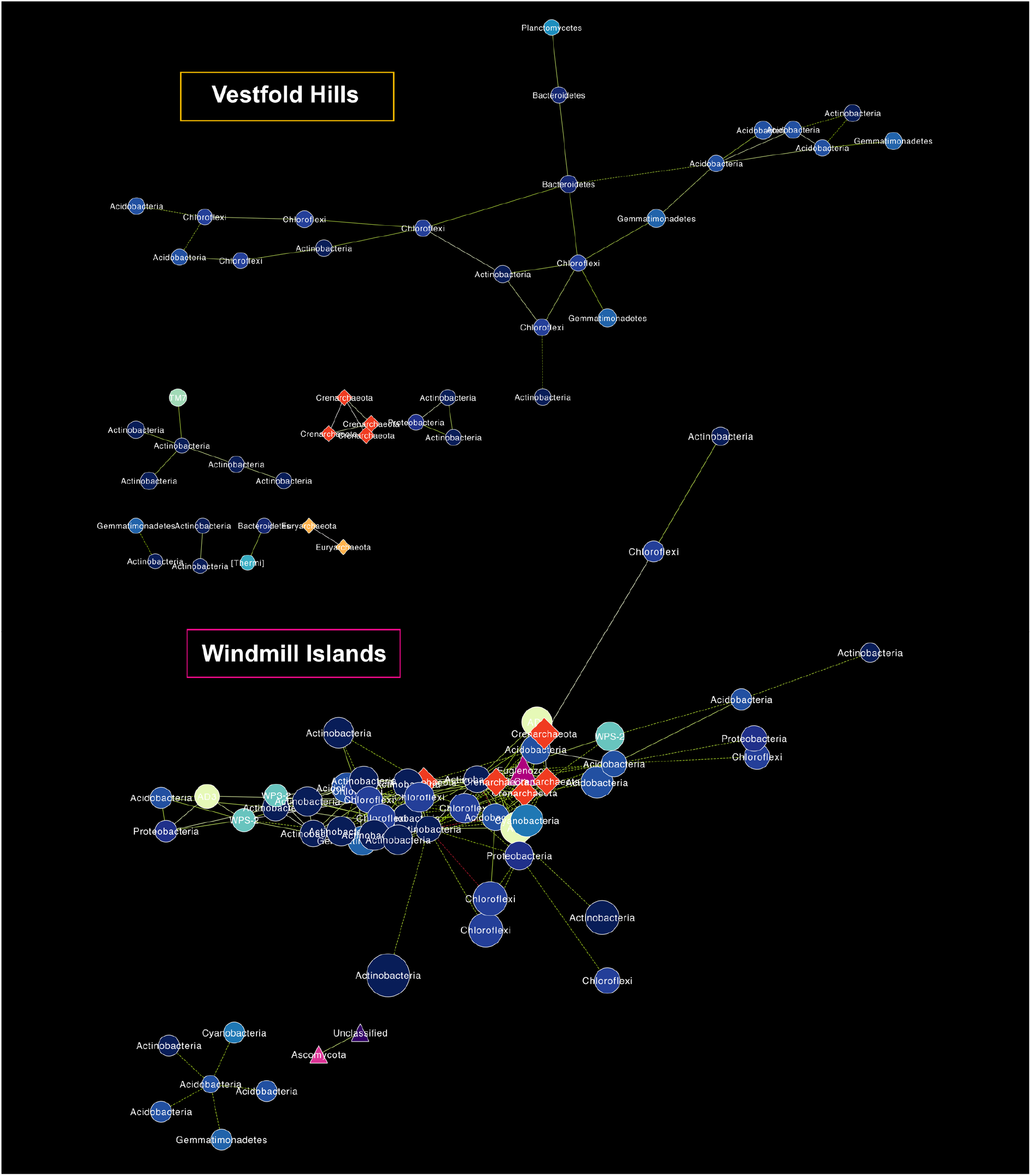
Domain-level OTU co-occurrence network of each OTU pair (*P*<0.001, *MIC*>0.8) across samples between the Windmill Islands and Vestfold Hills regions. Nodes (circles=bacteria, triangles=eukarya, diamonds=archaea) and edges are representative of individual OTUs and their correlations, respectively. Node size is proportional to their degree of connectivity and edge colour is based on linearity (green=positive, red=negative). Our soil microbial networks are comprised of moderately connected OTUs, more so at the Windmill Islands, structured amongst multiple components and forming a clustered topology. All three domains are present within the Windmill Islands network, whereas eukarya are absent from the Vestfold Hills network.

Co-occurrence OTU network analysis is a valuable exploratory tool to help ascertain potential biotic interactions, functional roles or identify ecological niches occupied by uncultured microorganisms in complex datasets (Barberán et al., 2012; Faust & Raes 2012; Ferrari et al., 2016). However, we acknowledge that significant co-occurrence patterns may not always have ecological relevance. Throughout both the Windmill Islands and Vestfold Hills, co-occurrence OTU patterns revealed interesting associations and potential sharing of niche space amongst many understudied taxa (Fig. 4). For instance, *Crenarchaeota* were prevalent in both regions but more associations at the Windmill Islands suggests different life histories or niche preferences between the two regions. Likewise, rare candidate bacterial phyla *Candidatus Eremiobactereota (WPS-2)* and *Candidatus Dormibactereota (AD3)* implicated in a novel mode of primary production termed trace gas chemosynthesis (Ji et al., 2017) only formed strong visible associations within the Windmill Islands network. Notable associations within the Vestfold Hills network included positive associations between *Saccharibacteria (TM7)*, a parasitic bacterium and *Actinobacteria* (Winsley et al., 2014). Also noted was a lack of co-occurrent eukaryotic species, suggesting competition. The astounding taxonomic diversity of *Actinobacteria* (Fig. 2), was reflected in their ability to occupy multiple niches and form the majority of connections with co-existing species, essentially moulding the microbial backbone within these Antarctic desert soils.

Information on non-random co-occurrence patterns are valuable for soil ecosystems where basic ecology and life history strategies of resident microbiota are largely unknown (Barberán et al., 2012; Janssen et al., 2006). This gap in knowledge is more apparent in extreme edaphic environments like Antarctica. Often described as the last great wilderness, Antarctica is the most undisturbed natural environment left on Earth (Terauds et al., 2012). But, the predicted acceleration of rapid ice-melt leading to large-scale biotic homogenisation raises many concerns regarding the potential for rare biodiversity loss (Terauds et al., 2017). We provide comprehensive information of the structuring of microbial biodiversity in east Antarctic soils. These findings provide a new understanding of the basic ecological concepts underlying Antarctic species abundance and distributions. By doing so, we now have a mechanistic basis for predicting potential effects of environmental disturbance at the micro-biodiversity scale.

## Materials and Methods

### Study area, soil sampling and physiochemical analysis

Sampling was performed by the Australian Antarctic Division (AAD) across nine polar desert locations spanning two ice-free regions (the Windmill Islands and Vestfold Hills) within the vicinity of Casey (66°17’S, 110°45’E) and Davis (68°35’S, 77°58’E) research stations in Eastern Antarctica (Fig. 1). Four sites were chosen from the Windmill Islands region: Mitchell Peninsula (MP: 66°31’S, 110°59’E); Browning Peninsula (BP: 66°27’S, 110°32’E); Robinson Ridge (RR: 66°22’S, 110°35’E); and Herring Island (HI: 66°24’S, 110°39’E). Five sites were chosen from the Vestfold Hills region: Adams Flat (AF: 68°33’S, 78°1’E); Old Wallow (OW: 68°36’S, 77°58’E); Rookery Lake (RL: 68°36’S, 77°57’E); Heidemann Valley (HV: 68°35’S, 78°0’E); and The Ridge (TR: 68°54’S, 78°07’E). At each site, soil samples (n=93) were taken along three parallel transects following a geospatial design (Siciliano et al., 2014). All soils (n=837) included in this study have been previously submitted for extensive physiochemical analysis by the AAD and Bioplatforms Australia (Table S2).

### DNA extraction and Illumina amplicon sequencing

Soil samples were extracted and quantified in triplicate using the FASTDNA™ SPIN Kit for Soil (MP Biomedicals, Santa Ana, CA, US) and Qubit™ 4 Fluorometer (ThermoFisher Scientific, NSW, Australia) as described in van Dorst et al., 2014. Diluted DNA (1:10 using nuclease-free water) was submitted to the Ramaciotti Centre for Genomics (University of New South Wales, Sydney, Australia) for amplicon paired-end sequencing on the Illumina MiSeq platform (Illumina, California, US) with controls in accordance to protocols from the Biome of Australia Soil Environments (BASE) project by Bioplatforms Australia (Bissett et al., 2016). We targeted bacterial 16S rRNA (n=837 total), eukaryotic 18S rRNA (n=162 total) and archaeal A16S rRNA (n=162 total) genes using the following primer sets: 27F/519R (Lane 1991); 1391f /EukBr (Amaral-Zettler et al., 2009); and A2F/519 (Reysenbach et al., 1995).

### Open OTU picking, assignment and classification

We followed the UPARSE-OTU algorithm (Edgar 2013) endorsed by Bioplatforms Australia by directly employing USEARCH 32-bit v10.0.240 (Edgar 2010) and VSEARCH 64-bit v2.8.0 (Rognes et al., 2016). Sequences were quality filtered, trimmed and clustered de novo to pick OTUs at 97% identity, reads were then assigned to separate sample-by-OTU matrices for each amplicon (Table S1). OTUs were taxonomically classified against the SILVA v3.2.1 SSU rRNA database (Quast et al., 2013). Where applicable, new OTU matrices were merged with existing ones using the QIIME 2 (https://qiime2.org) feature-table merge option. These were rarefied using the qiime feature-table rarefy function to generate random subsamples (bacterial 16S=700k reads, eukaryotic 18S=23k reads, archaeal 16S=850k reads).

### Multivariate and statistical analyses in R

All multivariate and statistical analyses were carried out in the R environment (R Core Team 2018). Subsampled rarefaction curves (q=0) were generated using the iNEXT package (Hsieh et al., 2018). Non-metric multidimensional scaling (NMDS) ordinations (distance=euclidean and bray-curtis) and chao1 richness estimates were calculated in vegan v2.5-3 (Oksanen et al., 2018). The ggcorrplot v0.1.2 package (Kassambara 2018) was used to compute a matrix of pearson’s correlation p-values (method=pearson, sig.level=0.05) between chao1 richness estimates and selected soil environmental parameters (where *R*<0.4 is described as weak, 0.4>*R*<0.6 as moderate; and R>0.6 as strong correlations). Unless specified otherwise, all plots were visualised using a combination of ggplot2 v3.1.0 (Wickham 2016) and ggpubr v0.2 (Kassambara 2018).

### Domain-level co-occurrence OTU network from abundance data

OTUs representing less than 0.001% of the total relative abundance of the bacterial, eukaryotic and archaeal communities within a given region were combined for network analyses (Fig. 4). Correlations between the relative abundance of each OTU pair across samples were calculated using the maximal information coefficient (*MIC*) in the MINE software package (Reshef et al., 2011). After correction for multiple testing (Benjamini and Hochberg 1995), statistically significant (*P*<0.001) co-occurrence relationships between pairs of OTUs were uploaded into the CYTOSCAPE software (Shannon et al., 2013) to generate network diagrams, displaying only very strong associations (*MIC*>0.8). Nodes (circle=bacteria, triangle=eukarya, diamond=archaea) and edges are representative of individual OTUs and their correlation between multiple nodes, respectively. The size of each node is proportional to their degree of connectivity and coloured according to phylogeny (Fig. 2). Edge colour is based on the positive or negative sign linearity. Statistical inferences of network topology were calculated using the Network Analyser algorithm (treatment=undirected) in CYTOSCAPE (Table S2).

### PLN and NB fitted species abundance distribution curves

As described in Connolly et al., 2013, poisson-lognormal (PLN) and negative binomial (NB) models were fitted to our empirical data using maximum likelihood methods representing niche and neutral distributions, respectively. Pooled species abundances were fitted on regional and individual site levels then displayed on a logarithmic scale (Fig. 5, S4 and S5). We refrained from pooling data based on individual phyla, as datasets with a small number of abundance values provide very little information on the shape of the SAD (Connolly et al., 2013).

**Figure 5.**
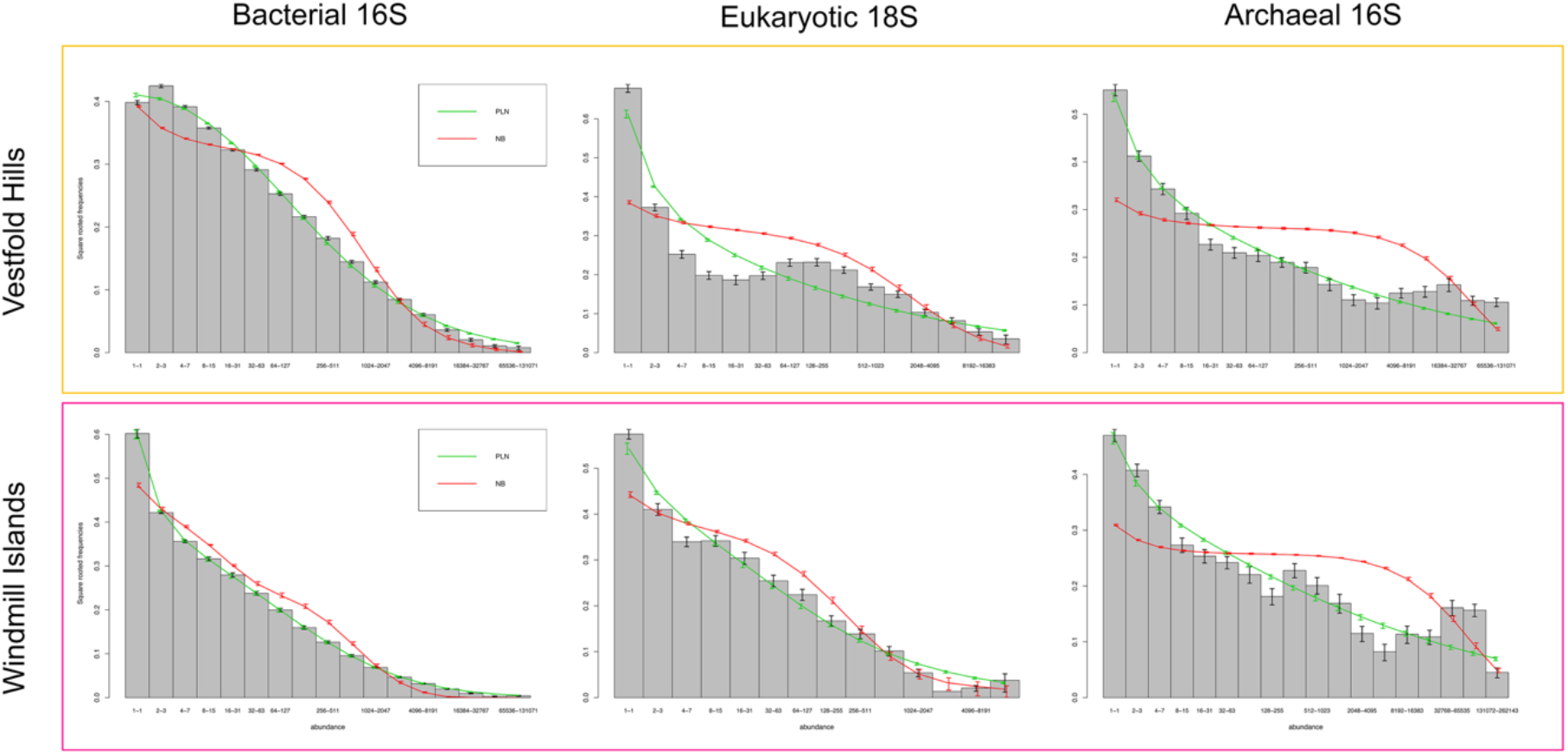
Fitted species abundance distribution (SAD) curves of polar soil microbial communities between the Vestfold Hills and Windmill Islands regions. The bars represent the mean proportion of species at each site in different octave classes of abundance. The green and red lines show the mean of fitted values from region-by-region fits of the poisson-lognormal (PLN) and negative binomial (NB) distributions to the data, respectively. A PLN-fit best explains the overall structure of these communities, particularly for bacterial communities at the Windmill Islands. Whereas, eukaryotic and archaeal communities across both regions exhibit multimodal distributions suggesting the emergence of neutrality.

### Deposition of data in an open source database

The datasets generated and analysed during the current study are all available through the Australian Antarctic Datacentre, [http://dx.doi.org/10.4225/15/526F42ADA05B1] and the BASE repository, [https://data.bioplatforms.com/organization/about/australian-microbiome].

## ACKNOWLEDGEMENTS

The authors would like to thank the Australian Antarctic Division for their logistical support in the successful collection of samples. We also thank Steven Siciliano for soil sampling, the Ramaciotti Centre for Genomics for their amplicon sequencing services and Bioplatforms Australia who supported provision of the Vestfold Hills biodiversity data.

Author contributions
BCF, MMT and EZ designed the study. AT co-ordinated sample collection and provided the environmental metadata. JvD, SW and EZ extracted the DNA for sequencing. EZ processed the sequencing data and performed the analyses. LMT provided scripts for the fitted species abundance distributions. EZ drafted the manuscript, and all authors read, collaborated, and approved the final manuscript.

