## Supplementary Figures and Tables for "Extreme niche partitioning promotes a remarkably high diversity of soil microbiomes across eastern Antarctica"

**This file contains**

Figures S1 to S4

Tables S1 to S3

**SUPPLEMENTARY FIGURES**

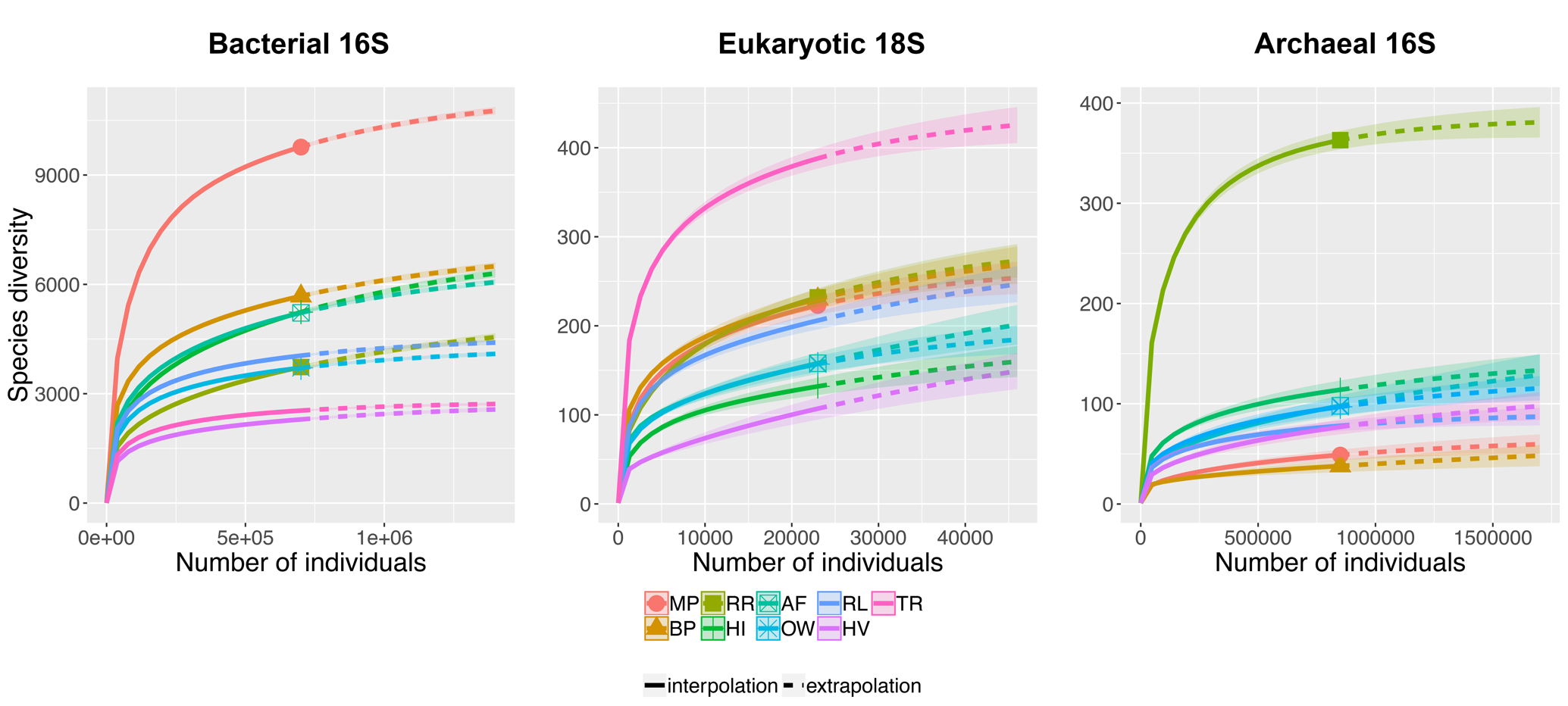

**Figure S1** Rarefaction curves of subsampled bacterial, eukaryotic and archaeal communities between sites. In all cases, asymptote was reached indicating that sufficient sampling depth had been achieved.

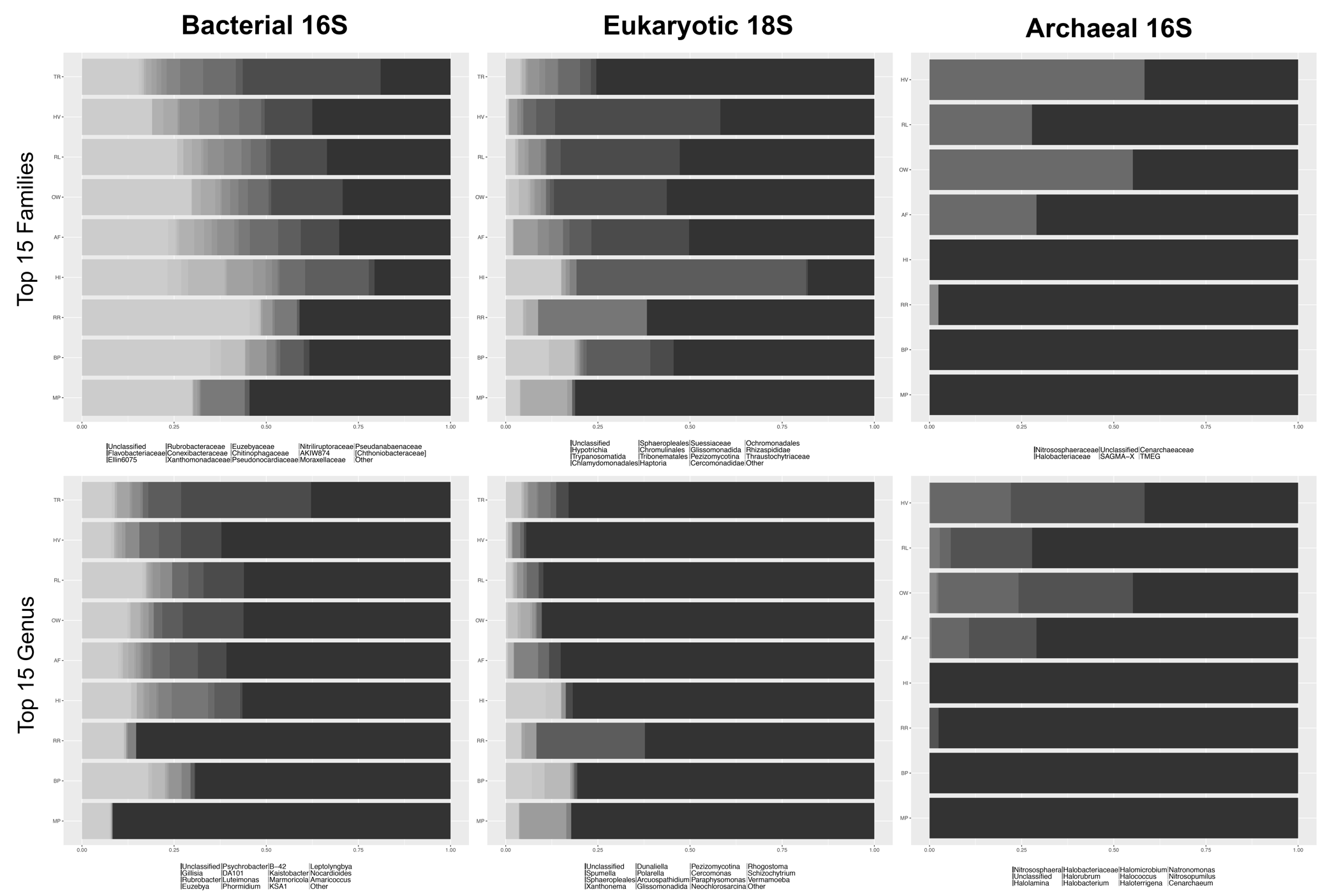

**Figure S2** Top 15 most abundant families and genus of bacterial, eukaryotic and archaeal communities between sites. As taxonomic levels decrease, the number of unclassified taxa increase substantially.

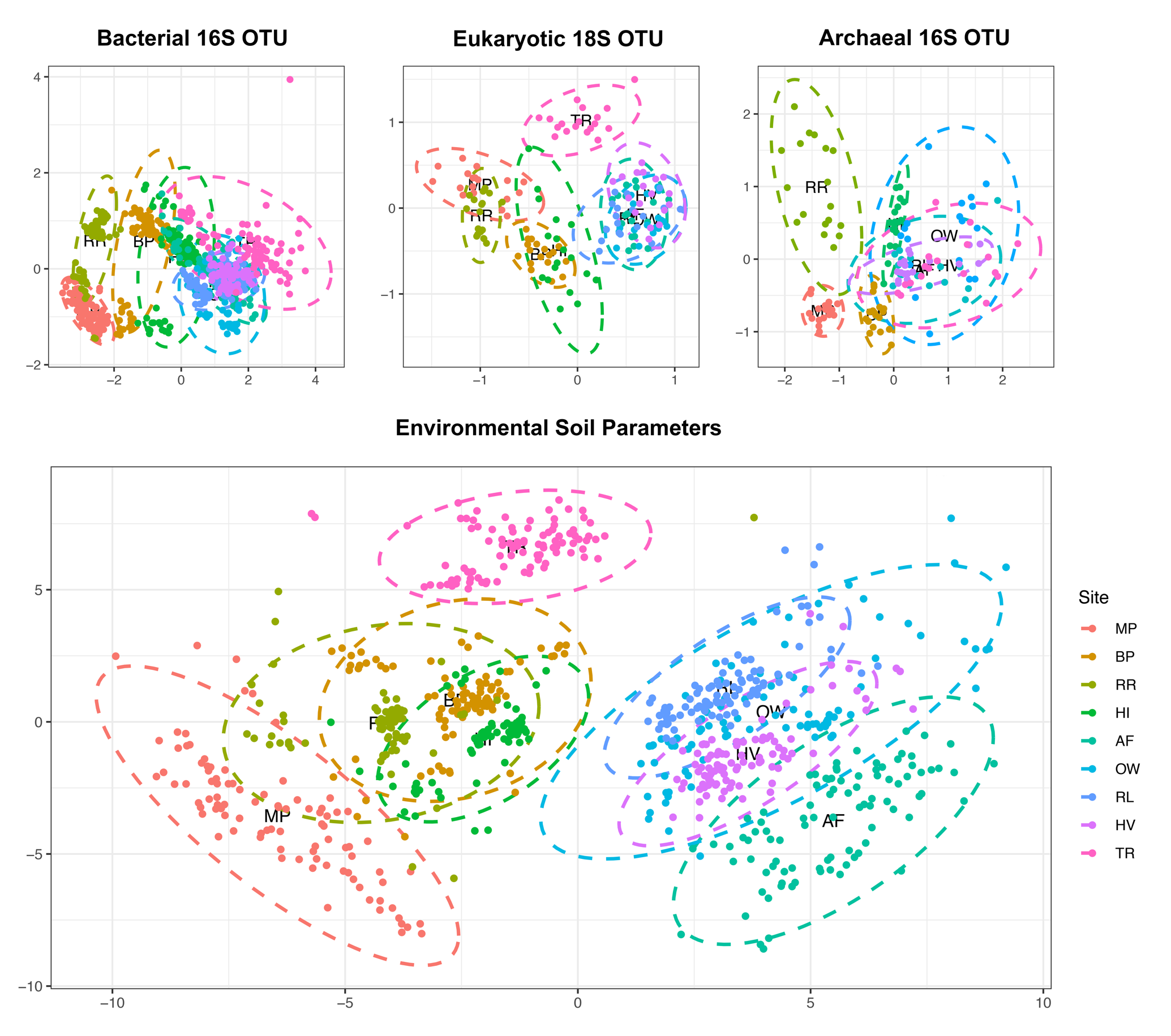
 **Figure S3** NMDS plots of microbial OTU communities and environmental soil datasets. All samples clustered according to site and broadly by geographic region.

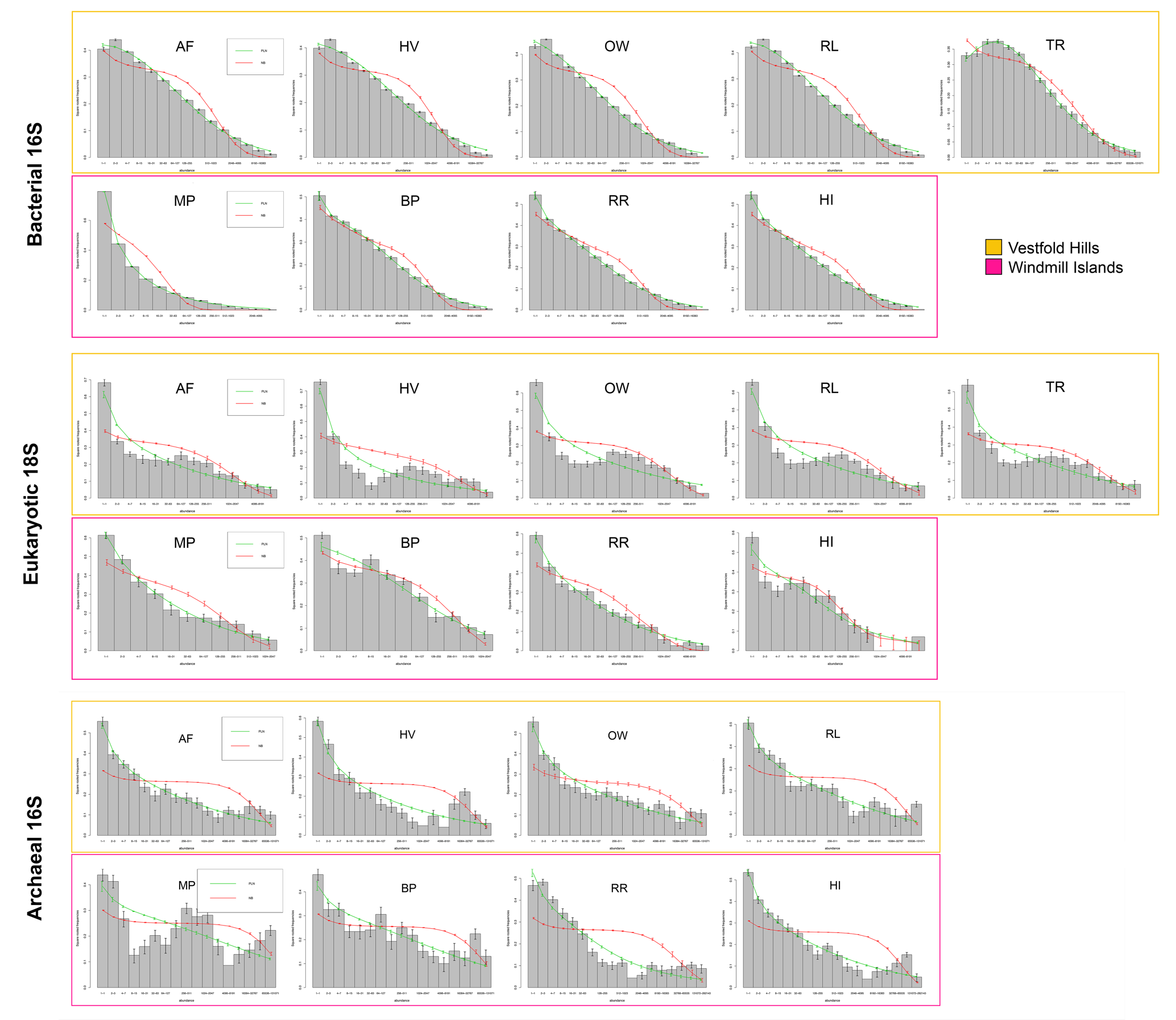

**Figure S4** Site level PLN and NB fitted SADs. Trends remain consistent with those observed for the region-level SADs (Fig. 5).

**SUPPLEMENTARY TABLES**

**Table S1** Summary of amplicon sequencing output and OTU pipeline analysis.

|  | **Samples per site** | **Total No. of Samples** | **Reads after Quality Filtering** | **OTUs** | **Singletons** |
| --- | --- | --- | --- | --- | --- |
| **Bacterial 16S** | 93 | 837 | 60,495,244 | 36,251 | 9,952 |
| **Eukaryotic 18S** | 18 | 162 | 1,299,519 | 1,511 | 329 |
| **Archaeal 16S** | 18 | 144 | 13,373,072 | 589 | 72 |

**Table S2** CYTOSCAPE network topology analysis between regions at the domain-level.

|  | **Vestfold Hills** | **Windmill Islands** |
| --- | --- | --- |
| **No. of Nodes** | 43 | 58 |
| **No. of Edges** | 44 | 201 |
| **Network Density** | 0.049 | 0.122 |
| **Network Heterogeneity** | 0.567 | 0.873 |
| **Clustering coefficient** | 0.214 | 0.448 |
| **Connected components** | 8 | 3 |
| **Network diameter** | 7 | 6 |
| **Network radius** | 1 | 1 |
| **Network centralisation** | 0.074 | 0.401 |
| **Shortest paths** | 518 (28%) | 2482 (75%) |
| **Characteristic path length** | 3.247 | 2.377 |
| **Average no. of neighbours** | 2.047 | 6.931 |

**Table S3** Environmental soil parameters averaged between sites.

|  | **Parameter** | **MP** | **BP** | **RR** | **HI** | **AF** | **OW** | **RL** | **HV** | **TR** |
| --- | --- | --- | --- | --- | --- | --- | --- | --- | --- | --- |
| **Geographical** | Elevation (m) | 31.76 | 41.24 | 35.67 | 30.92 | 3.01 | 18.9 | 12.13 | 11.68 | 27.91 |
|  | Slope (degree) | 9.48 | 2.8 | 6.91 | 6.94 | 0.72 | 1.94 | 3.24 | 1.95 | 5.69 |
|  | Aspect | 217.6 | 172.76 | 119.28 | 130.46 | 137.59 | 188.93 | 165.11 | 143.8 | 201.46 |
|  | DMF | 0.97 | 0.89 | 0.91 | 0.96 | 0.98 | 0.92 | 0.92 | 0.93 | 0.96 |
|  | Conductivity (uS/cm) | 49.54 | 36.62 | 23.39 | 163.03 | 2623.94 | 6361.04 | 3061.61 | 1738.61 | 2898.69 |
|  | pH | 5.41 | 6.57 | 5.3 | 6.72 | 7.92 | 8.43 | 7.96 | 8.85 | 8.87 |
| **Nutrients** | Total C (mg C/kg DMB) | 1475.84 | 1938.89 | 2586.56 | 691.67 | 1420.95 | 3942.67 | 1414.59 | 1713.72 | 774.28 |
|  | Total N (mg N/kg DMB) | 46.38 | 184.62 | 235.53 | 152.35 | 132.78 | 303.5 | 168.28 | 140.06 | 81.75 |
|  | Total P (mg P/kg DMB) | 28.48 | 800 | 40.42 | 1069.44 | 981.11 | 707.78 | 1655.56 | 729.44 | 753.26 |
| **Water Extractable Ions** | Cl Water (ppm) | 50.99 | 42.83 | 6.1 | 130.85 | 3899.05 | 4287.37 | 2603.72 | 1783.88 | 1964.78 |
|  | NO_2_ Water (ppm) | 0.08 | 0.08 | 0.13 | 0.08 | 0.76 | 0.31 | 0.54 | 0.7 | 1 |
|  | Br Water (ppm) | 0.08 | 0.11 | 0.14 | 0.19 | 5.34 | 11.3 | 2.42 | 0.95 | NA |
|  | NO_3_ Water (ppm) | 0.35 | 1.75 | 0.67 | 0.87 | 15.1 | 7.06 | 21.94 | 3.95 | 6.09 |
|  | PO_4_ Water (ppm) | 0.63 | 1.17 | 4.65 | 5.06 | 3.86 | 2.07 | 5.04 | 3.71 | 2.06 |
|  | SO_4_ Water (ppm) | 14.15 | 9.57 | 6.94 | 29.76 | 1079.66 | 1489.86 | 375.28 | 634.11 | 1549.56 |
| **Elemental Cation Levels** | P (mg/kg) | 22.43 | 12.17 | 45.11 | 64.56 | 9.12 | 8.96 | 36.09 | 18.54 | NA |
|  | K (mg/kg) | 0.49 | 54.85 | 0.86 | 94.07 | 135.7 | 241.59 | 418.82 | 205.52 | 268.45 |
|  | Ca (mg/kg) | 0.16 | 114.62 | 0.1 | 94.59 | 971.97 | 3283.52 | 842.45 | 1171.69 | 965.67 |
|  | Mg (mg/kg) | 28.17 | 116.92 | 70.31 | 55.13 | 303.92 | 1348.22 | 476.13 | 449.35 | 268.45 |
|  | Zn (mg/kg) | 2.32 | 1.19 | 1.79 | 0.96 | 1.17 | 1.93 | 1.59 | 1.59 | 1.01 |
|  | B (mg/kg) | 136.92 | 0.31 | 377.52 | 0.51 | 4.28 | 4.92 | 5.01 | 5.42 | 9.59 |
|  | S (mg/kg) | 1.28 | 10.02 | 4.82 | 17.86 | 498.17 | 1393.84 | 471.57 | 490.22 | 648.35 |
|  | Cu (mg/kg) | 50.54 | 2.25 | 25.42 | 2.57 | 4.95 | 11.4 | 7.17 | 8.76 | 3.9 |
|  | Fe (mg/kg) | 328.74 | 137.45 | 518.8 | 132.85 | 267.12 | 240.69 | 219.4 | 195.69 | 152.04 |
|  | Mn (mg/kg) | 3.11 | 17.92 | 4.37 | 2.3 | 7.58 | 34.04 | 43.33 | 20.36 | 19.56 |
|  | Na (mg/kg) | 0.59 | 54.22 | 1.07 | 239.31 | 2457.62 | 5124.22 | 3348.4 | 1630.17 | 2148.84 |
|  | Al (mg/kg) | 0.1 | 267.26 | 0.45 | 184.68 | 102.83 | 184.47 | 199.33 | 183.9 | 135.92 |
|  | CECe (meq/100g) | 0.07 | 1.92 | 0.1 | 2.21 | 40.79 | 72.79 | 46.16 | 39.56 | 16.63 |
|  | Ca (meq/100g) | 0.22 | 0.57 | 0.11 | 0.47 | 4.85 | 16.38 | 4.2 | 5.85 | 4.83 |
|  | Mg (meq/100g) | 0.02 | 0.97 | 0.04 | 0.45 | 2.5 | 11.1 | 3.92 | 3.7 | 1.73 |
|  | K (meq/100g) | 16.55 | 0.14 | 37.71 | 0.24 | 0.35 | 0.62 | 1.07 | 0.53 | 0.69 |
|  | Na (meq/100g) | 31.77 | 0.24 | 35.48 | 1.04 | 10.69 | 22.29 | 14.56 | 7.09 | 9.34 |
|  | %CEC Ca | 38.01 | 25.3 | 15.04 | 18.27 | 8.86 | 15.49 | 7.46 | 12.84 | 26.06 |
|  | %CEC Mg | 484.44 | 42.49 | 881.11 | 19.08 | 4.84 | 12.93 | 8.02 | 8.98 | 15.12 |
|  | %CEC K | 608.33 | 6.2 | 1550 | 10.07 | 0.75 | 0.83 | 2.11 | 1.39 | 6.58 |
|  | %CEC Na | 4050.5 | 10.35 | 9672.22 | 39.9 | 20.22 | 23.21 | 25.45 | 15.94 | 49.37 |
| **Particle Size** | Mud % | 1.03 | 1.51 | 1.49 | 1.26 | 7.14 | 24.93 | 23.37 | 25.86 | NA |
|  | Sand % | 65.52 | 71.04 | 69.36 | 69 | 84 | 67.36 | 64.31 | 61.86 | NA |
|  | Gravel % | 32.37 | 23.18 | 25.18 | 28.14 | 8.86 | 7.71 | 12.32 | 12.29 | NA |
|  | Minimum (um) | 47.09 | 17.35 | 24.94 | 35.97 | 13.94 | 3.04 | 2.96 | 2.22 | NA |
|  | Maximum (um) | 1248.34 | 1185.89 | 1120.16 | 1193.39 | 1104.52 | 834.81 | 989.32 | 905.01 | NA |
|  | Mean (µm) | 465.04 | 410.21 | 349.29 | 440.94 | 321.54 | 211.32 | 213.49 | 198.67 | NA |
|  | Quartile Deviation (PQD) | 1.06 | 1.14 | 1.39 | 1.06 | 1.16 | 1.86 | 1.84 | 1.84 | NA |
|  | Sorting Coeff (So) | 0.48 | 0.45 | 0.39 | 0.48 | 0.45 | 0.28 | 0.29 | 0.28 | NA |
|  | Graphic Skewness | -0.21 | -0.21 | -0.18 | -0.24 | -0.18 | -0.24 | -0.08 | -0.1 | NA |
|  | IGS | -0.25 | -0.31 | -0.22 | -0.29 | -0.24 | -0.28 | -0.14 | -0.16 | NA |
|  | Kurtosis | 0.95 | 1.16 | 0.92 | 0.99 | 1.19 | 0.94 | 0.98 | 0.98 | NA |
| **Oxide Levels** | SiO_2_ (%) | 33.32 | 31.67 | 35.56 | 36.55 | 62.68 | 60.41 | 57.84 | 61.35 | NA |
|  | TiO_2_ (%) | 0.61 | 0.93 | 0.98 | 0.93 | 0.8 | 0.84 | 1.07 | 0.86 | 60.15 |
|  | Al_2_O_3_ (%) | 85.61 | 86.7 | 86.32 | 85.88 | 13.81 | 13.1 | 14.8 | 13.45 | 0.92 |
|  | Fe_2_O_3_ (%) | 6.6 | 5.89 | 6.27 | 6.89 | 7.85 | 9.45 | 9.73 | 9.42 | 14.24 |
|  | MnO (%) | 0.25 | 0.11 | 0.13 | 0.14 | 0.12 | 0.14 | 0.12 | 0.14 | 9.19 |
|  | MgO (%) | 1.86 | 1.39 | 1.84 | 3.04 | 4.08 | 4.78 | 4.47 | 4.7 | 0.12 |
|  | CaO (%) | 2.39 | 3.3 | 3.88 | 5.3 | 5.33 | 5.24 | 5.32 | 5.25 | 4.45 |
|  | Na_2_O (%) | 2.78 | 2.81 | 2.84 | 2.95 | 3.56 | 3.22 | 3.89 | 2.98 | 5.06 |
|  | K_2_O (%) | 3.14 | 3.08 | 3.1 | 2.88 | 1.47 | 1.55 | 2.28 | 1.55 | 3.64 |
|  | P_2_O_5_ (%) | 0.2 | 0.2 | 0.38 | 0.26 | 0.21 | 0.19 | 0.39 | 0.19 | 1.92 |
|  | SO_3_ (%) | 0.05 | 0.01 | 0.04 | 0.06 | 0.17 | 0.4 | 0.22 | 0.15 | 0.24 |
|  | Cl (ppm) | 66.03 | 95.33 | 123.25 | 395.06 | 5463 | 13164.28 | 6528.61 | 2962.22 | 0.22 |
